# A framework for estimating the determinants of spatial and temporal variation in vital rates and inferring the occurrence of unobserved extreme events

**DOI:** 10.1101/033720

**Authors:** Simone Vincenzi, Dušan Jesenšek, Alain J Crivelli

## Abstract

We develop an overarching framework that combines long-term tag-recapture data and powerful statistical and modeling techniques to investigate how population, environmental, and climate factors determine variation in vital rates and population dynamics in an animal species, using as a model system the population of brown trout living in Upper Volaja (Western Slovenia). This population has been monitored since 2004; Upper Volaja is also a sink, receiving individuals from a source population living above a waterfall. We estimate the numerical contribution of the source population on the sink population and test the effects of temperature, population density, and extreme events on variation in vital rates among more than 2,500 individually tagged brown trout. We found that fish dispersing downstream from the source population help maintain high population densities in the sink population despite poor recruitment. The best model of survival for individuals older than juveniles includes additive effects of year-of-birth and time. Fast growth of older cohorts and higher population densities in 2004-2005 suggest very low population densities in late1990s, which we hypothesize were caused by a flash flood that strongly reduced population size and created the habitat conditions for faster growth and transient higher population densities after the extreme event.

## 1 Introduction

The way that vital rates (e.g., growth, survival, movement, fecundity), life-history traits (e.g., age and size at maturity, lifespan), and life histories (the timing of key events in an organism’s lifetime and the trade-offs between life-history traits) in a population or species change in space and time and the consequences of this variation for population dynamics are central topics in ecology and evolutionary biology [1]. With unprecedented rates of climate (e.g., mean and variance of temperature and precipitation) [2] and environmental (e.g., habitat fragmentation, pollution) change [3], we need to develop and integrate powerful methods for *(i)* estimating of growth, survival, reproductive traits, and movement in animal populations and *(ii)* testing theory-based hypothesis on the relationship between populations and environment that can help us forecast future population size, age-, size-, and spatial structure to inform species management [4].

Due to the large number of potential determinants and individual and group heterogeneity in responses, understanding how variation in habitat and population factors drives variation in traits and population dynamics is intrinsically difficult. This task is further complicated by small population sizes, demographic stochasticity, and the occurrence of stochastic – and potentially unobserved – environmental and climate events strongly affecting vital rates, population size, and population dynamics [5]. For instance, many flash floods with dramatic effects on population go unobserved or unreported, although their occurrence can be inferred from compensatory changes in vital rates that can be captured by novel, powerful statistical models [6,7]. In addition, most monitoring programs follow either only a fraction of the population under investigation or the population is part of a meta-population or source-sink system. In those cases, estimates of vital rates and life-history traits can be misleading when movement is not explicitly considered, since movement within populations and dispersal among populations can introduce substantial bias [8].

Within a population, habitat factors - both extrinsic (e.g., weather, predators or food availability) and intrinsic (e.g., population density or composition) - and their interaction [9], determine a large part of the temporal variation in the distribution of vital rates, recruitment, and population dynamics [10]. Part of the variation is often explained by individual heterogeneity, since in many taxa organisms living in the same population greatly differ in the ability to acquire resources, in their life-history strategies, and in their contribution to the next generation [11]. These differences may result from complex interactions between genetic, environmental, population factors, and chance, and often have substantial consequences for both the ecological and evolutionary dynamics of species [12]. When not accounted for, the presence of individual variation can bias the estimation of vital rates and other demographic traits for use in population and life-history models [6,7], which may translate to incorrect inference on co-variation among vital rates and timing of life-history events, and inaccurate predictions of those models [13].

In addition to the intrinsic biological and computational complexities associated with the estimation of variation in vital rates, taking individual heterogeneity into account increases the complexity of model specification and parameter estimation [6,14]. Longitudinal data (e.g., tag-recapture) and random-effects models greatly facilitate the estimation of individual and group (i.e. sex, year-of-birth cohort) variation in vital rates, life-history traits, and fitness [15].

Our goal is a to provide a overarching framework for the fine-grained estimation of vital rates and their determinants – including extreme climate events - when there is substantial individual and shared variation in those traits, using a freshwater salmonid population as a model system. Specifically, we combined a long-term tag-recapture dataset and an overarching statistical and modeling framework to estimate variation in vital rates among years, groups, and individuals of a population of brown trout *Salmo trutta* L. living in Upper Volaja (Western Slovenia). Then, we tested hypotheses on the determinants of variation to understand how that variation influences the population dynamics of the species and its future under scenarios of climate change.

The brown trout population of Upper Volaja is enclosed between two impassable waterfalls and receives fish from a un-sampled population living upstream; it is thus a “source-sink system”, in which the dynamics of the population of Upper Volaja depends not solely on its internal demography, but also on the input of individuals from the source population living above the waterfall [16].

Specifically, we used our framework to estimate and test hypotheses on (1) the contribution of the source population to the sink population, (2) the effects of water temperature, population density, location within stream, and extreme climate events on variation in vital rates (growth, survival, movement, recruitment, i.e. the main axes of variation) among more than 2,500 individually tagged fish that have been sampled between 2004 and 2015.

## 2 Material and Methods

### 2.1 Species and study area

The population of Upper Volaja was initiated in the 1920s by stocking brown trout *Salmo trutta* L., with no additional stocking since then. The monitored population of Upper Volaja lives in a stretch of stream approximately 265 m in length that is enclosed between two impassable waterfalls (Lat: 46.229386 N, Long: 13.659794 E; Table ESM 1). The catchment is pristine and there is neither poaching nor angling in the stream. Brown trout is the only fish species living in Upper Volaja. Fish can disperse from the upstream part of the population (a source population for Upper Volaja) into Upper Volaja and from Upper Volaja into the downstream population (which is thus a sink for the upstream population(s)). Due to the harsh environment (steep, slippery rocks), sampling has never been conducted above the waterfall (AW henceforth) or below (BW) the waterfalls enclosing Upper Volaja, although we know that the population of brown trout in AW extends for ∼400 m. Within Upper Volaja, there are no physical barriers impairing upstream or downstream movement of brown trout; however, the movement of brown trout throughout their lifetime is typically limited [17].

#### 2.1.1 Sampling

We sampled the population of Upper Volaja bi-annually in June and September each year from September 2004 to September 2015 (23 sampling occasions). Fish were captured by electrofishing and length (*L*) and weight (*W*) recorded to the nearest mm and g, respectively. If captured fish had *L* > 115 mm, and had not been previously tagged or had lost a previously applied tag, they received an external Carlin tag [18] and age was determined by reading scales. Carlin tags were assigned to a total of 2,647 fish. Fish are aged as 0+ in the first calendar year of life, 1+ in the second year and so on. Sub-yearlings are smaller than 115 mm in June and September, so fish were tagged when at least aged 1+. The adipose fin was also removed from all fish captured for the first time (starting at age 0+ in September), including those not tagged due to small size at age 1+. Therefore, fish with intact adipose fin were not sampled at previous sampling occasions at age 0+ or 1+. Fish were also assigned a sampling location (sector) within Upper Volaja. Sectors were numbered from 4 (most upstream) to 1, with sector 4 being the longest (95 m) and sector 1 the shortest (45 m) (Table ESM 1).

#### 2.1.2 Environmental data

Annual rainfall recorded between 1985 and 2013 in the meteorological station closest to Upper Volaja (Lat: 46.25 N, Long: 13.58333 E, Kobarid, Slovenia) ranged between 1600 and 3400 mm. The maximum daily rainfall over the same time period was recorded on November 7^th^ 1997 (252 mm) (Fig. ESM 1). An ONSET temperature logger recorded mean daily water temperature in Upper Volaja. Missing water temperature data in 2004 (all year) and 2005 (from January 1^st^ to June 10^th^) were estimated using water temperature recorded in the stream Lipovscek (Pearson’s *r* = 0.98 for daily water temperature data between 2004 to 2014). Annual growing degree-days *(GDDs,* [19]) and mean annual *T* showed very little variation from 2004 to 2014 (mean±sd *GDDs* = 1204.94± 115.71, CV = 9%; *T* = 8.37± 0.21, CV = 3%) (Fig. ESM 2). Water flow rates have never been recorded in Upper Volaja. A full list of abbreviations used in this paper is in Table ESM 2.

### 2.2 Density and movement

We estimated density of 0+ fish only in September, since fish emerged a few days before the June sampling. We estimated density of fish older than 0+ for age, size-class, or cohort using a two-pass removal protocol [20] as implemented in the R [21] package FSA [22]. Three passes provided the same results as two passes [23]. Total stream surface area (746.27 m^2^) was used in the estimation of fish density (in fish ha^-1^). We assessed the contribution of trout from AW (source) to Upper Volaja (sink) by estimating for each year of sampling the proportion of fish that were not sampled in Upper Volaja either at 0+ in September or 1+ in June. Fish with adipose fin cut were assumed to be born in Upper Volaja or be early incomers (i.e., fish migrating into Upper Volaja when younger than 1+ in September), while fish with intact adipose fin were assumed to be born in AW and be “late incomers”, that is fish dispersing into Upper Volaja when 1+ in September or older. We grouped together fish born in Upper Volaja and early incomers (both “early incomers” from now on), since we cannot distinguish between them (see Text ESM 1 for methodological details). We tested for recruitment-driven population dynamics by estimating correlations between density of 0+ fish (*D*_0+_) in September and density of older than 0+ (*D*_>0+_) one or two years later.

For movement, we first estimated the proportion of tagged trout sampled in different sectors at different sampling occasions. Then, we estimated the parameters of a Generalized Linear Model (GLM) in which the number of different years in which a fish was sampled predicted the probability of a fish being sampled in different sectors.

### 2.3 Growth and body size

In order to characterize size-at-age and growth trajectories, we modeled (a) variation in size at first sampling (i.e., 0+ in September), (b) individual, year-of-birth cohort, and spatial variation in lifetime growth trajectories, and (c) variation in growth between sampling occasions.

#### 2.3.1 Variation in size at age 0+

We used linear regression to model the variation in mean length of cohorts at age 0+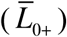 using *D*_>0+_and *GDDs* (up to August 31^st^) and their interaction as candidate predictors. We log-transformed both *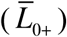*and *D*_>0+_ [10]. We carried out model selection with the MuMIn package [24] for R, using the Akaike Information Criterion (AIC) as a measure of model fit. We considered that models had equal explanatory power when they differed by fewer than 2 AIC points [25].

#### 2.3.2 Lifetime growth trajectories

The standard von Bertalanffy Growth Function (vBGF) is

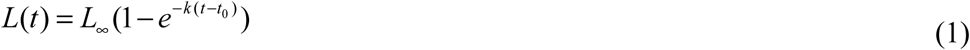

where *L*__∞__ is the asymptotic size, *k* is a coefficient of growth (in time^-1^), and *t*_0_ is the (hypothetical) age at which length is equal to 0.

In the vast majority of applications of the vBGF, *L*_∞_, *k*, and *t*_0_ have been estimated at the population level starting from cross-sectional data, without accounting for individual heterogeneity in growth. However, when data include measurements on individuals that have been sampled multiple times, failing to account for individual variation in growth will lead to biased estimations of mean length-at-age [6,7].

We used the formulation of the vBGF specific for longitudinal data of [6], in which *L*_∞_ and *k* may be allowed to be a function of shared predictors and individual random effects. In the estimation procedure, we used a log-link function for *k* and *L*_∞_, since both parameters must be non-negative. We set

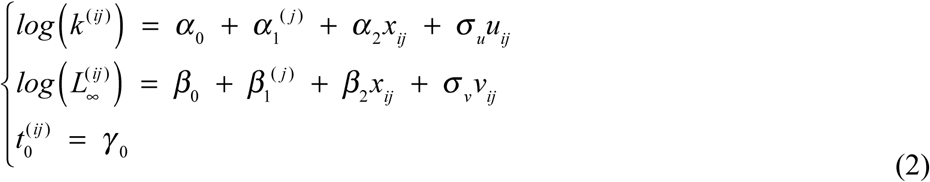

where *u* ∼ *N* (0,1) and *v* ∼ *N* (0,1) are the standardized individual random effects, *σ*_*u*_ and *σ*_*v*_ are the standard deviations of the statistical distributions of the random effects, *i* is the individual, *j* is the index for groups (e.g. cohort), and the other parameters are defined as in Eq. (1). The continuous predictor *x*_*ij*_ (i.e. population density or temperature, as explained below) in Eq. (2) must be static (i.e. its value does not change throughout the lifetime of individuals).

Models were fitted with the Automatic Differentiation Model Builder (ADMB), an open source statistical software package for fitting nonlinear statistical models [26,27]. One of the features of ADMB is the ability to fit generic random-effects models using an EB approach using the so-called Laplace approximation [28]. Unlike most mixed model software available, ADMB is totally flexible in its model formulation, allowing any likelihood function to be coded up in C++. The flexibility is useful if the model involves several individual-specific parameters. The gradient (i.e. the vector of partial derivatives of the objective function with respect to the parameters) provides a measure of convergence of the parameter estimation procedure in ADMB. Although speed consideration and model complexity may motivate the use of a less strict convergence criterion, by default ADMB stops when the maximum gradient component is < 10 ^-4^.

Since the growth model operates on an annual time scale and more data on tagged fish were generally available in September of each year, we used September data for modeling lifetime growth. Following [6] and [7], we included three potential predictors of *k* and *L*_∞_ : (*i*) cohort (*Cohort*) as a group (i.e. categorical) variable (*α*_1_ and *β*_1_ in Eq. 2), (*ii*) population density (fish older than 0+) in the first year of life (*D*_>0 +,*born*_) as a continuous variable (i.e. *x*_*ij*_ Eq. 2), and (*iii*) *GDDs* in the first year of life as a continuous variable.

In addition, we tested the hypothesis of a longitudinal gradient in growth within Upper Volaja-as commonly found in marble trout *Salmo marmoratus* living in the study area [6] - in which fish living more upstream show higher length-at-age than fish living more downstream, probably due to more food drift available to them. Thus, we also used (*iv*) sampling sector as categorical predictor of *k* and *L*_∞_. Following [6], *Cohort* and sampling sector were introduced as fixed effects. Datasets for the analysis of lifetime growth trajectories are described in Text ESM 2.

#### 2.3.3 Growth in size between sampling intervals

We used Generalized Additive Mixed Models (GAMMs) [29] to model variation in mean daily growth *Gd* (in mm d-^1^) between sampling occasions using length *L*, *Age*, *GDDs* over sampling intervals by *Season*, and *D*_>0+_ as predictors, plus fish ID as a random effect. Since we expected potential non-linear relationships between the two predictors and *G*_*d*_, we used candidate smooth functions for *L* and *GDDs*. We carried out model fitting using the R package *mgcv* [30] and model selection as in the Section *“Variation in size at age 0+”.*

### 2.4 Recruitment

Brown trout living in Upper Volaja spawn in December-January and offspring emerge in June-July. Females achieve sexual maturity when bigger than 150 mm, usually at age 2+ or older, and can be iteroparous [31,32]. We used density of fish with *L* > 150 mm as density of potential spawners at year *t* (*D*_s,t_) and density of 0+ in September of year *t* as a measure of recruitment (*R*_t_).

The most popular stock-recruitment models (e.g. Ricker’s, Beverton-Holt’s, Cushing’s) rarely provide a good fit to recruitment data of small freshwater fish populations. Following [23], we thus used Generalized Additive Models (GAMs) to model variation in *R*_t_ using density of potential spawners in September of year *t*-1 (*D*_s,t-1_) and *GDDs* for year *t* up to emergence time (we assumed from January 1^st^ to May 31^st^ for standardization purposes) as predictors. We used candidate smooth functions for *GDDs* and *D*_s,t-1_ as we were expecting non-linear relationships between the two predictors and *R*_t_. We carried out model selection as in Section *“Variation in size at age 0+”*.

### 2.5 Survival

To characterize variation in survival and identify the determinants of this variation, we modeled survival between sampling occasions for tagged fish and survival between age 0+ and 1+ for untagged fish that were first sampled in the stream when 0+.

#### 2.5.1 Survival of tagged individuals

Our goal was to investigate the effects of mean temperature, early density, season, age, and sampling occasion on variation in probability of survival of tagged fish using continuous covariates (*D*_>0+_, mean temperature between sampling intervals *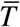*, *Age*) at the same time of categorical predictors (*Cohort*, *Time*, *Season*). Since only trout with *L* > 115 mm (aged at least 1+) were tagged, capture histories were generated only for those fish. Full details of the survival analysis are presented in Text ESM 3.

Two probabilities can be estimated from a capture history matrix: *ϕ*, the probability of apparent survival (defined “apparent” because it includes permanent emigration from the study area, which is basically inevitable in mobile species when only a fraction of the area occupied by the species is studied), and *p*, the probability that an individual is captured when alive [15]. We used the Cormack–Jolly–Seber (CJS) model as a starting point for the analyses [15]. We started with the global model, i.e. the model with the maximum parameterization for categorical predictors. From the global model, recapture probability was modeled first. The recapture model with the lowest AIC was then used to model survival probabilities.

We modeled the seasonal effect (*Season*) as a simplification of full time variation, by dividing the year into two periods: June to September (*Summer*), and the time period between September and June (*Winter*). Since length of the two intervals (*Summer* and *Winter*) was different (3 months and 9 months), we estimated probability of apparent survival on a common annual scale. Both *Age* and *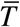* were introduced as either non-linear (as B-splines) or linear predictors, while *D*_>0+_ was introduced only as a linear predictor. In addition, we tested whether probability of apparent survival of trout that was born in AW was different from that of fish born in Upper Volaja. In this case, we used a subset of the whole dataset that included cohorts born between 2004 and 2010. We carried out the analysis of probability of survival using the package *marked* [14] for R.

#### 2.5.2 Survival from age 0+ to 1+ (first overwinter survival)

Because fish were not tagged when smaller than 115 mm (thus 0+ were never tagged as they are always smaller than 115 mm), we assumed a binomial process for estimating the probability *σ*_0+_ of first overwinter apparent survival (0+ in September to 1+ in June) for trout that were sampled in September of the first year of life and had the adipose fin cut (see Text ESM 4 for details on the estimation of *σ*_0+_). This way, immigration of un-sampled individuals at 0+ would not bias the estimates of apparent survival probabilities. We tested for density-dependent apparent survival *σ*_0+_ by estimating a linear model with *D*_>0 +,*m*_ (mean of *D*_>0+_ at year *t* in September and *t*+1 in June) as predictor of the estimate of 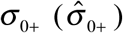. We log-transformed both 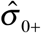_+_ and *D*_>0 +,*m*_. We carried out model selection as in Section *“Variation in size at age 0+”*.

## 3 Results

### 3.1 Variation in density, recruitment, and movement

The estimated probability of capture at every depletion pass was high (mean± sd of point estimates across sampling occasions: 0.86± 0.07 for 0+ fish and 0.91± 0.02 for fish older than 0+) (Table ESM 3). Population density was variable through time, although the coefficient of variation (CV) was low for *D*_>0+_ (15%) and high for *D*_0+_ (65%) (Fig. 1 and Table ESM 3). The estimated number of trout in the stream (mean± se) was between 0± 0 (year 2014) and 65± 1 (2015, 871± 19 fish ha^-1^) for 0+ and between 327± 2 (2015, 4382± 25 fish ha^-1^) and 548± 3 (2004, 7343± 38 fish ha^-1^) for older fish (Fig. 1 and Table ESM 3).

**Fig. 1.**
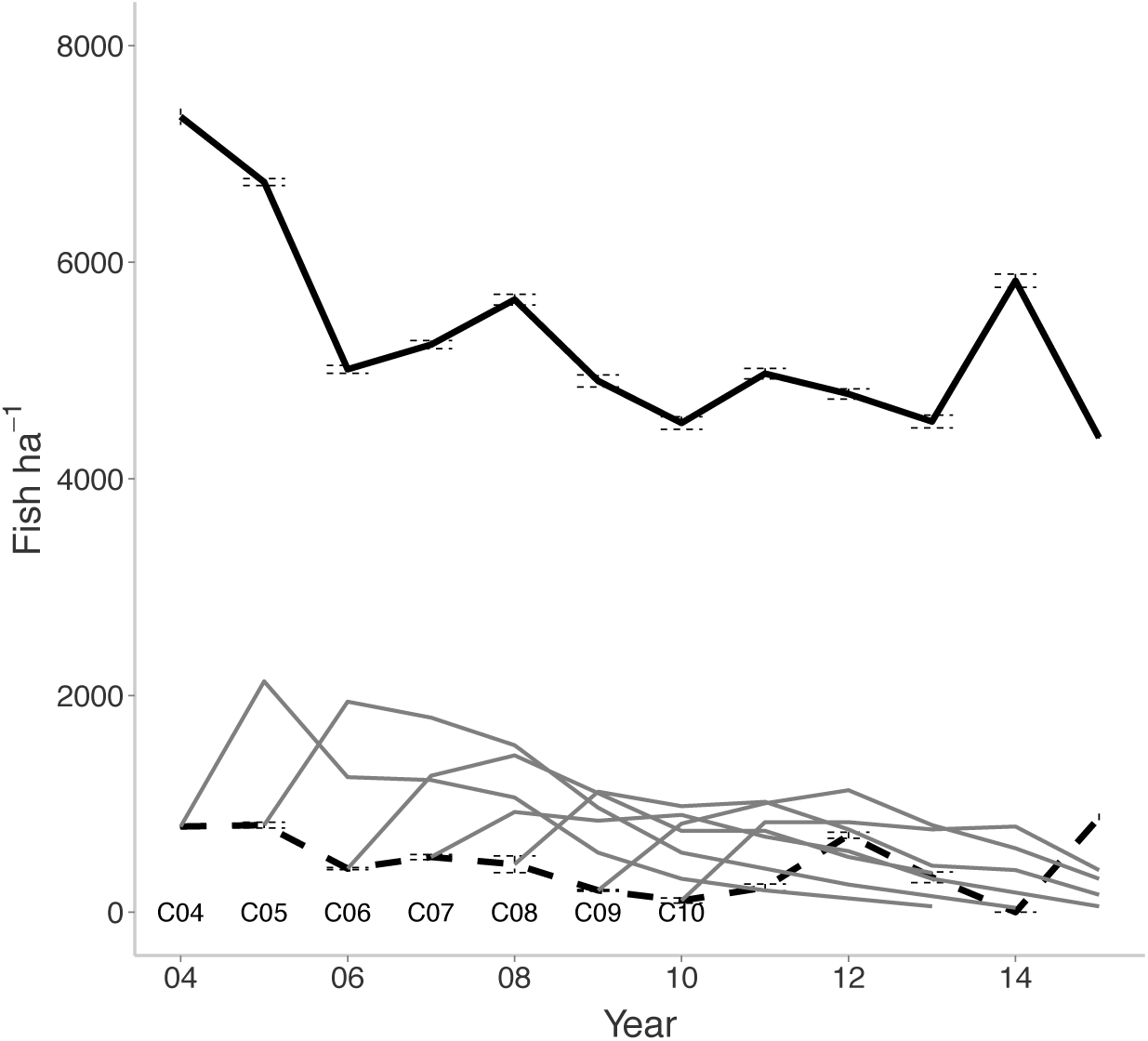
Density over time of brown trout aged 0+ (dashed black line), older than 0+ (solid black line), and single year-of-birth cohorts (from C04 = 2004 to C10 = 2010) in September. 95% confidence intervals are barely visible as the probability of fish capture at each passage was very high (∼90%) and consequently confidence intervals very narrow.

For each cohort born after the start of sampling (i.e., since 2004), the number of sampled fish in a cohort increased after the first sampling (i.e., from 0+ to 1+), thus showing that fish from AW contributed to population size and population dynamics of brown trout in Upper Volaja (Fig. 1). Since 2010 (the first year in which fish from cohorts born before 2004 were fewer than 10% of population size), the proportion of trout alive that were not sampled in Upper Volaja early in life (i.e., “late incomers”) has been high and stable across years (0.35± 0.05, Table ESM 4).

There was little variation in density of potential spawners across years (mean± sd = 3459± 442 fish ha^-1^, CV = 13%) and the best model of recruitment *R*_t_ did not include either *GDDs* or *D*_s,t-1_. We observed complete a recruitment failure in 2014, despite an average density of potential spawners sampled in September 2013 (3270± 21 fish ha^-1^). The estimated number of 1+ in September 2015 (all fish coming from AW after September 2014) was 20± 0.8.

There was no significant lagged correlation (either for lag of 1 or 2 years) between *D*_0+_ and *D*_>0+_, which indicates that recruitment (density of 0+ in September at year *t*) was not driving variation in population density of fish older than juveniles at year *t* + 1 or *t* + 2.

Only 26± 1% of tagged fish were sampled in more than one sector across sampling occasions. Of those, ∼25% were sampled at different sampling occasions in non-adjacent sectors, thus most movement was relatively limited in distance. The probability of being sampled in different sectors increased with the number of years in which a fish was sampled (GLM: *α* =-0.04± 0.02, *β* = 0.13± 0.01, *p*<0.01).

### 3.2 Growth and recruitment

The best model for mean length of age 0+ fish 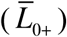 in September had only *D*_>0+_ as predictor (negative effect, *R*^2^ = 0.28, *p* = 0.06), although models with only *GDDs* (positive effect of *GDDs*, ΔAIC from best model = 0.61) or no predictors (ΔAIC = 0.78) had the same explanatory power of the best model.

Empirical growth trajectories for tagged fish (i.e., *L* > 115 mm) in September (*n* of length data = 4590; *n* of individuals in cohorts between 7 [cohort 2000] and 370 [2003]) showed individual variation in growth rates and size-at-age (Fig. 2 and Fig. ESM 3), thus supporting the choice of a growth model with individual random effects. The best growth model for brown trout had *Cohort* as a predictor of both *L*__∞__ or *k* (Table ESM 5), although the effect size of the difference in growth among cohort was small, in particular for cohorts born after 2003 (Fig. 2 and Table ESM 6). The biggest brown trout sampled in Upper Volaja was had *L* = 297 mm when 7 years old (Fig. ESM 3).

**Fig. 2.**
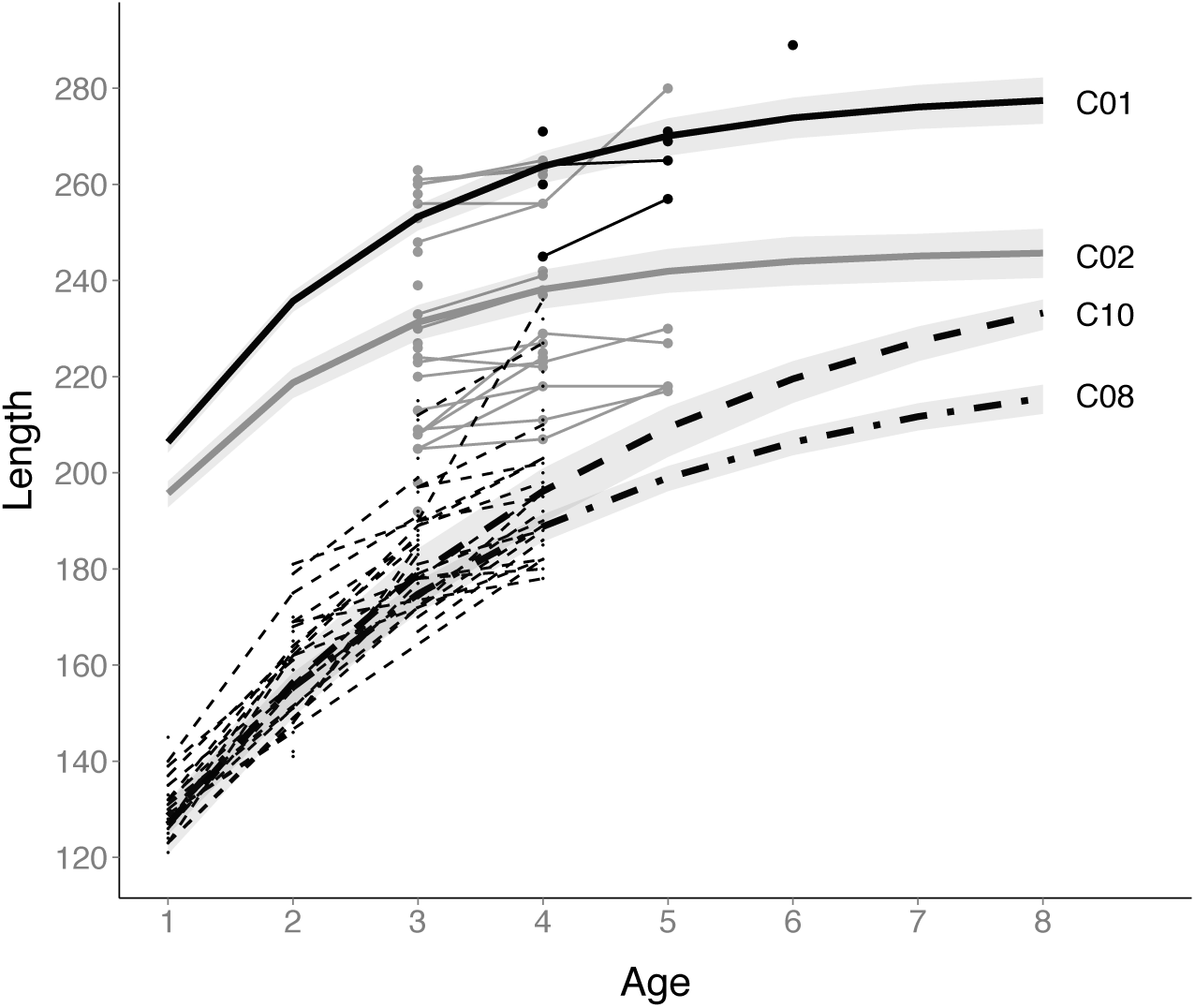
Average growth trajectories of brown trout (individual random effects for *L*_∞_ and *k* set to 0) living in different cohorts along with 95% confidence intervals of the average trajectories. Thin lines and dots are growth trajectories and length-at-age data of fish born in 2000 (black solid), 2001 (gray solid), 2010 (black dashed). We hypothesize that the bigger average size of fish born in 2000 and 2001 was a compensatory response to very low densities in late 1990s/early 2000s, probably caused by an extreme climate event.

We found a longitudinal gradient in lifetime growth trajectories in Upper Volaja, with fish sampled in sampling sectors more upstream growing faster and having larger asymptotic size than fish sampled in sectors more downstream. However, differences in mean length-at-age were small and confidence intervals for the average sector-specific growth trajectories tended to overlap (Fig. 3 and Table ESM 7).

**Fig. 3.**
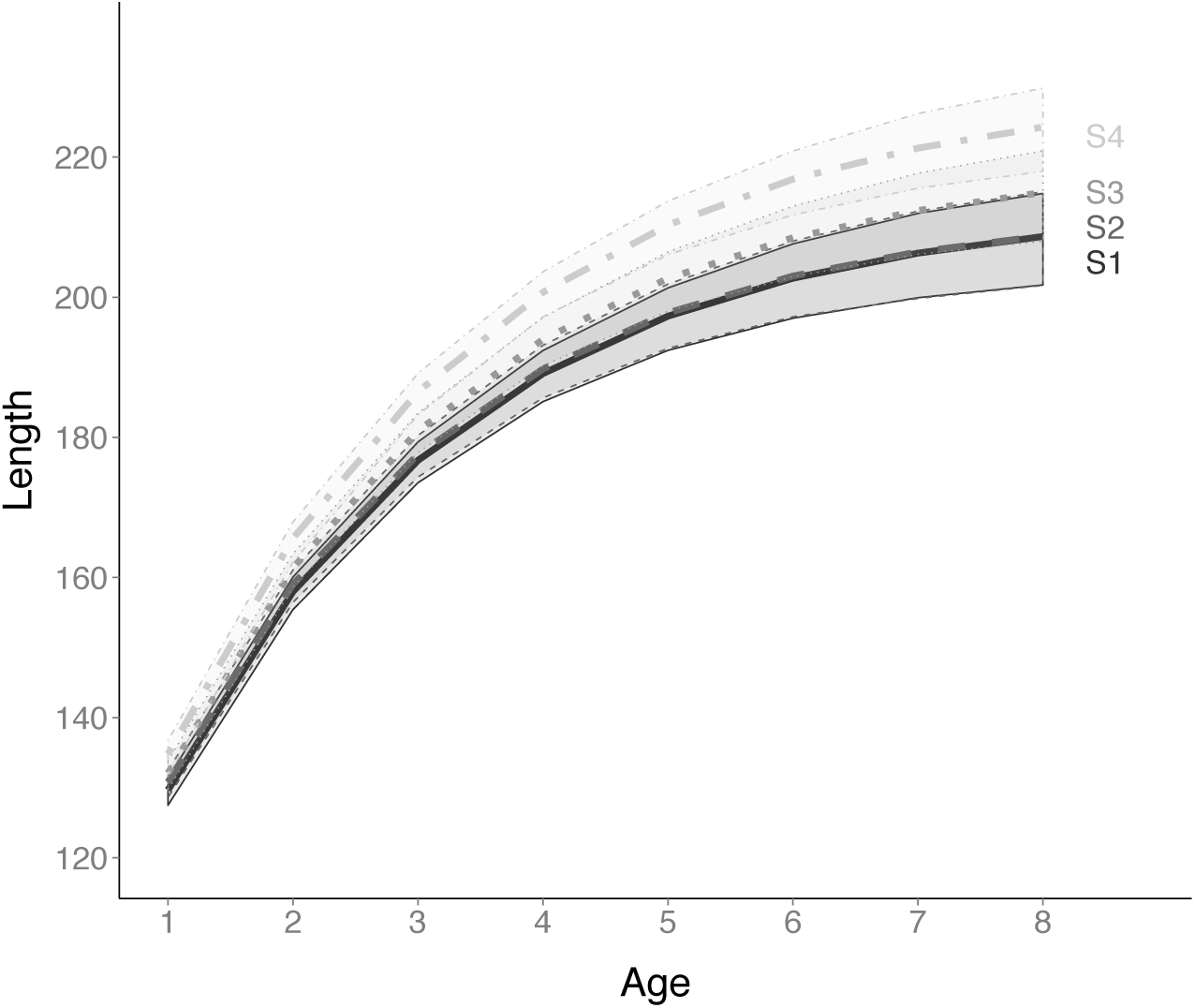
Average growth trajectories in sampling sectors for brown trout that have been sampled either once at age 1+ or multiple times in the same sampling sector (with first sampling occurring at age 1+). Brown trout tend to grow faster in sampling sectors more upstream (S4 is the most the upstream sampling and S1 the most downstream).

Brown trout relatively large early in life tended to remain larger than their conspecifics throughout their lifetime (Pearson’s *r* of size at age 1+ and size at age 3+ = 0.25, *p* < 0.01). The best model of growth between sampling intervals included *Cohort*, *Age* (growth tended to be slower at older ages), *L* (growth decreased with increasing *L*), and the interaction between density and *Season* as predictors (*n* = 4174, *R*^2^ = 0.33; Fig. ESM 4). *GDDs* had a positive, although small, effect on *Summer* growth and a negative and stronger effect on *Winter* growth (Fig. ESM 4).

### 3.3 Survival

We found a highly variable probability of early apparent survival *σ*_0+_ over years, ranging from 0.1 to 0.62 on an annual temporal scale. The best model for *σ*_0+_ included only *D*_>0 +,*m*_ as predictor, but contrary to what is commonly observed, the effect of *D*_>0 +,*m*_ on*σ*_0+_ was **positive** (linear model on log-log scale: *α* =-24.18± 11.06, *β* = 2.70± 1.30, *&&&*, *p* = 0.07).

Probability of capture for tagged brown trout was high (*p* = 0.84 on average across sampling occasions), with variation in probability of capture best explained by sampling occasion (Table ESM 8). The probability of apparent survival *ϕ* varied across sampling occasions, it was consistently greater than 0.4 on an annual temporal scale, and in some sampling occasions close to 0.8 (Fig. 4b, average survival (mean[95%CI]): 0.55[0.54-0.57]). The probability of survival of fish born in Upper Volaja and “early incomers” and “late incomers” from AW was basically the same (“early incomers” – mean[95%CI]: 0.56[0.55-0.59]; “late incomers”: 0.60[0.57-0.63]). The best model for probability of apparent survival had an additive effect of *Cohort* and *Time* on *ϕ* (Table 1 and Fig. 4a), but part of the variation in *ϕ* due to *Cohort* may be explained by *Age* (Fig. ESM 5). Population density had small effects on *ϕ*, with a slight tendency toward lower *ϕ* at higher densities (Fig. 4c).

**Fig. 4.**
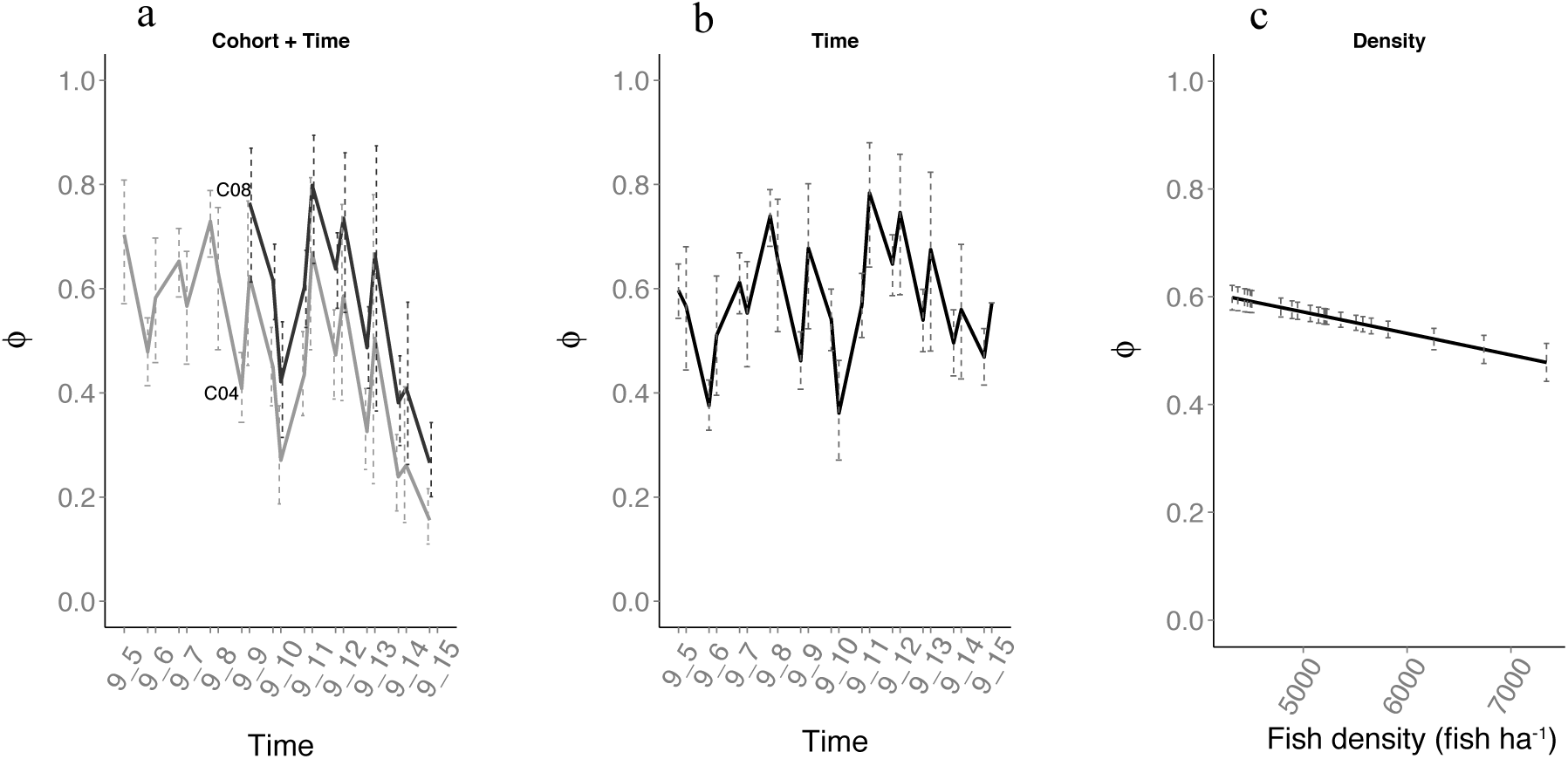
Probability of survival (mean and 95% confidence intervals) on an annual temporal scale with the model with additive effect between *Cohort* and *Time* (best model, panel a), model with *Time* effect (b), and model with Density of fish older than 0+ (*D*_>0+_) effect (c).

**Table 1.** Best 10 models for the probability of survival *ϕ* using time-varying probability of capture (i.e. *p*(*Time*)). The symbol * denotes interaction between predictors. *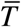* is mean temperature between sampling occasions; *D*_>0+_ is density of fish older than 0+; *Time* = interval between two consecutive sampling occasions; *Season* is a categorical variable for *Summer* (June to September) and *Winter* (September to June). *bs* means that the relationship temperature and probability of survival has been modeled as a B-spline function. *npar* = number of parameters of the survival model.

| Model | npar | AIC | $\Delta AIC$ | weight |
| --- | --- | --- | --- | --- |
| $\phi(Cohort + Time) p(Time)$ | 58 | 13279.56 | 0.00 | 1.00 |
| $\phi(Cohort * Age) p(Time)$ | 52 | 13382.15 | 102.59 | 0.00 |
| $\phi(Cohort + Age) p(Time)$ | 38 | 13406.20 | 126.64 | 0.00 |
| $\phi(Time) p(Time)$ | 44 | 13432.99 | 153.43 | 0.00 |
| $\phi(Season * Age) p(Time)$ | 26 | 13435.53 | 155.97 | 0.00 |
| $\phi(bs(Age)) p(Time)$ | 26 | 13437.87 | 158.31 | 0.00 |
| $\phi(Season + Age) p(Time)$ | 25 | 13439.00 | 159.44 | 0.00 |
| $\phi(Age) p(Time)$ | 24 | 13447.41 | 167.85 | 0.00 |
| $\phi(Cohort + Season) p(Time)$ | 38 | 13515.85 | 236.29 | 0.00 |
| $\phi(Cohort) p(Time)$ | 37 | 13518.33 | 238.77 | 0.00 |

## 4 Discussion

In order to understand how variation in vital rates and life histories of organisms emerge we need (a) long-term studies that include contrasting environmental conditions [5], (b) longitudinal data [15], and (c) statistical models that can tease apart shared, and individual contributions to the observed temporal and spatial variation in vital rates, life histories, and population dynamics [33]. Our overarching statistical and modeling framework integrated all three components, allowing us to quantify variation in density, dispersal, movement, growth, recruitment, and survival, and provide clear answers to some of the hypotheses tested on the determinants of temporal and spatial variation in vital rates and population processes.

Population density in Upper Volaja was very stable after the first 3 years of sampling and seemed to be unaffected by variation in recruitment. This occurred because the large proportion of fish living in Upper Volaja that were born above the waterfall (>30% of population size) was largely buffering variation in recruitment. Average growth trajectories of cohorts were very similar to each other except for the two older cohorts, which were characterized by much faster growth than that of fish born in later years. High population densities in 2004-2005 and fast growth of fish born in early 2000s point to very low population densities in late 1990s/early 2000s, probably a consequence of an extreme climate event (e.g., flash flood or debris flow) that caused high mortalities. We did not find any strong effect of either temperature or population density on vital rates of brown trout living in Upper Volaja, probably due low variation in both density of brown trout and water temperature since 2004.

We discuss how the framework we presented can facilitate the integration of population-level processes across temporal and spatial scales and identify gaps that can be informed by an understanding of those processes. We also discuss what we have learned about the population of Upper Volaja, the pieces of missing information that would further our understanding of demographic and life-history processes in freshwater salmonids, and how our results help advance our understanding of those processes in natural populations.

### 4.1 Growth and extreme events

The best model of brown trout lifetime growth trajectories included year-of-birth cohort as a categorical predictor for both *L*_∞_ and *k*. The vBGF parameters can seldom be interpreted separately, especially when only a few older fish are measured [6]; it follows that the analysis of the whole growth trajectories is necessary for understanding growth variation among individuals and cohorts. The method for estimating lifetime growth trajectories we used in this work provides excellent predictions of unobserved size-at-age [6,7]. We found that most of the differences among the average growth trajectories of cohorts were due to fish born in years 2000 and 2001, which grew much faster than fish born in later years. We did not observe evident effects of density on either lifetime growth or growth between sampling occasions in Upper Volaja for fish born after 2003, likely because year-to-year variation in density in Upper Volaja since 2004 was too small to induce noticeable effects on growth [5].

We hypothesize that very low population densities in late 1990s/early2000s created the conditions for (i) faster growth of fish born in early 2000s and (ii) higher population densities in 2004 and 2005, and likely since 2000. The most likely explanation for the very low population densities in late 1990s/early2000s is an extreme climate event – a flash flood or a debris flow-that caused massive fish mortalities. As found in other salmonids [34], the relaxation of density-dependent pressure and fewer older fish occupying profitable stream habitat may allow brown trout to grow faster and have higher-than-average survival and production of young, the latter leading to transient higher population densities. [23,32] found an increase in individual growth, survival, recruitment, and population density between 3 and 4 years after flash floods affecting the Slovenian marble trout populations of Lipovscek and Zakojska. The increases in growth and population density in marble trout were comparable to those observed in the brown trout population of Upper Volaja [32]. In addition, in marble trout there was no recruitment in the two years following the flash flood [32] and since 2004 in Upper Volaja we did not sample any brown trout born before 2000. Our hypothesis is also supported by rainfall data; an extreme rainfall event was recorded in the rainfall station closest to Upper Volaja (Kobarid) in 1997 (252 mm of rainfall on November 7^th^, 95^th^ percentile of 1961-2013 daily rainfall maxima), with rainfall similar to that recorded near Lipovesck (303.5 mm) and Zakojska (223.5 mm) when flash floods occurred in December 2007 [32]. Finally, in the monitored population of marble trout closest to Upper Volaja (Zadlascica, Lat: 46.22437 N, Long: 13.77999 E), population density estimated in 1998 (the first year of sampling) and after the flood of 2007 that affected many marble trout populations were basically the same (∼120 adult fish ha-^1^), with a rapid increase in population density observed in late 1990s/early 2000s [23].

Daily rainfalls similar to that of 1997 were recorded in December 25^th^ 2009 (247 mm) and November 5^th^ 2012 (235 mm). In those days, floods were recorded in several Western Slovenian streams [32], but in Upper Volaja heavy rainfalls were not followed by population collapses, noticeable lower survival rates, or visible alterations of stream morphology. Flash floods are usually generated by short-period (typically a few hours) intense rainfall (minimum rainfall for the formation of flash floods varies with local geography) on small, steep catchments that exceeds drainage capacity in urban areas or infiltration capacity in rural areas [35]; colluvium transported by water - ranging in size from fine-grained material to wood to large boulders – strongly increases the lethal effects of flash floods on fish. It follows that heavy rainfall is necessary, but not sufficient for flash flood formation (topography, soil conditions, and ground cover also play important roles), that flash floods may have vastly different effects on fish depending on the material transported by water, and that daily rainfall may only be correlated with probability of flash flood formation. Nevertheless, the extreme daily rainfall maximum recorded in December 2009 and the extreme annual rainfall recorded in 2010 (3374 mm, the highest since 1961) may have been among the determinants of the lower-than-average apparent survival of brown trout estimated in 2009 and 2010, for instance through a high number of fish displaced downstream.

Along with higher estimated densities, recorded extreme rainfall, and compensatory responses similar to those observed in other salmonids, the random-effects vBGF was crucial for developing the robust hypothesis of the occurrence of an extreme climate events causing massive mortality in the late 1990s [6]. In fact, using only the few data points at older ages that were available for the older cohorts would not allow estimating their cohort-specific average lifetime growth trajectories [6]. Similarly, the estimation of a correlation structure among extreme rainfall events in Western Slovenia that leverages information from tens of meteorological stations [36] and future measurement of water flows in small streams with probes or meters would provide a clearer picture of the past and future extreme rainfall events and flash floods in Western Slovenian streams. This would also help us interpret some currently unexplained observations in Upper Volaja that may depend on climate events, such as recruitment failures.

#### 4.2 Recruitment and movement

Whether there is a relationship between the number or density of spawners (i.e., stock) and recruitment in freshwater fishes has been a subject of debate for decades, and contrasting results have been found. For instance, a Ricker stock-recruitment relationship was found in 5 populations of brown trout living at the periphery of its distribution (Spain, Nicola et al. 2008), although egg production and density of the spawning stock were not observed, but estimated from fecundity, trout density, and proportion of sexually mature trout. On the contrary, stock-recruitment relationships were not found in brown trout living in 4 sites within Rio Chaballos (also in Spain), where environmental factors – in particular flow rates – were found to mostly determine recruitment [38]. We did not find any evidence of a relationship between potential spawners and recruitment in Upper Volaja, although the density of potential spawners estimated according to size is only a crude proxy of the density of the actual spawners [32]. In addition, due the influx of fish from upstream, we cannot exclude that young-of-the-year – despite suitable spawning areas in Upper Volaja – were in part or largely produced above the waterfall.

In Upper Volaja, we found that number of fish for some cohorts increased through multiple years, indicating that large numbers of older fish were dispersing from the source population into the sink population. In addition, 26% of tagged fish (thus older than 0+) within Upper Volaja changed sector at least once throughout their lifetime, although those movements may be of just tens of meters over a lifetime. Contrary to what we found for marble trout living in the area [23], the population size of brown trout older than juveniles was not driven by recruitment, a result that should be mostly ascribed to the large proportion of fish older than young-of-the-year that were born above the waterfall.

#### 4.3 Survival

Although density-dependent early survival is commonly found in brown trout [10], there are examples of brown trout populations showing density-dependent survival only at the adult stage [39] or constant loss rates [40]. In the brown trout population of Upper Volaja, probability of survival early in life was some years lower than and in other years comparable to probability of survival of older fish (i.e. between 0.1 and 0.62 annual survival probability). It is often challenging to compare survival probabilities reported in the literature for natural populations, since more intuitive temporal scales (i.e., month or year) are not always used, and the delta method [41] or similar approaches must be used for estimating the standard errors of the transformed survival probabilities.

In Norwegian streams with similar water temperature and life-history traits to the Upper Volaja population, first overwinter survival was at lower end of the range found for Upper Volaja (0.65 to 0.87 monthly survival for the 9 “winter” months, i.e. ∼ between 0.01 and 0.20 annual survival) [42]. Density appeared to have had a positive effect on first overwinter survival in Upper Volaja, although the relationship was noisy and more years of data are needed to be more confident on the effects of density on early survival. One hypothesis for a transient positive correlation between density and survival is favorable environmental conditions (e.g., abundant food, optimal water flow) in Upper Volaja and upstream leading to higher production and then higher dispersal from upstream to Upper Volaja and higher survival in Upper Volaja.

Probability of survival of tagged fish in Upper Volaja between sampling occasions was quite variable, with no evidence of a winter demographic bottleneck. Variable survival had minor effects population density, since the influx of fish from AW seems to buffer the effects of variation in survival. Neither water temperature nor population density seemed to explain much variation in survival, probably due to the little variation observed since 2004 for both temperature and density; variation in survival might thus be ascribed to variation in flow rates or other unobserved properties of the environment. For instance, water flows in winter or spring that were strong enough to displace eggs, kill larvae before emergence, or damage spawning grounds - but not to cause higher-than-average mortalities among fish - may have caused the recruitment failure observed in 2014. Other years of very low recruitment may be similarly explained by particularly high water flows in winter or spring; for instance, in a brown trout population living in a Austrian Alpine river (Ybbs River), it was found that high water flows during incubation and emergence were negatively correlated with recruitment success [43].

Fish from AW seem to have only a “numeric” effect on the population of Upper Volaja, although fish genotyping and molecular pedigree reconstruction [32,44] will allow testing hypotheses on the effects of place of birth of fish on recruitment, whether the Upper Volaja population would be self-sustaining without the steady input of brown trout from upstream, and the occurrence of recent genetic bottlenecks caused by extreme climate events [32]. Survival probabilities in Upper Volaja for fish older than juveniles were greater than in Norwegian streams with similar water temperature and populations of brown trout with life-history traits comparable to those of the Upper Volaja population (winter: ∼0.25; summer: ∼0.48, [45]). “Real” survival might be substantially higher than “apparent” survival, thus indicating that Upper Volaja provides a very favorable habitat for brown trout. In fact, the large number of fish migrating from AW leaves open the possibility that many fish permanently left Upper Volaja to disperse into the population living below the waterfall, thus leading to a substantial underestimation of “real” survival.

## Acknowledgements

We thank the employees and members of the Tolmin Angling Association (Slovenia) for carrying out fieldwork since 1993.

## Data Accessibility Statement

Tag-recapture data - fig**share**: http://dx.doi.org/10.6084/m9.figshare.1617848

## Ethics/Fieldwork permissions statement

All sampling work was approved by the Ministry of Agriculture, Forestry and Food of Republic of Slovenia and the Fisheries Research Institute of Slovenia. Sampling was supervised by the Tolmin Angling Association (Slovenia).

## Authors Contribution Statement

SV conceived the ideas and designed methodology; AJC conceived and run the Marble trout Project and AJC and DJ collected the data; SV analyzed the data; SV, AJC, and DJ led the writing of the manuscript. All authors gave final approval for publication.

